# Statistical mechanics of phase space partitioning in large-scale spiking neuron circuits

**DOI:** 10.1101/132993

**Authors:** Maximilian Puelma Touzel, Fred Wolf

## Abstract

Synaptic interactions structure the phase space of the dynamics of neural circuits and constrain neural computation. Understanding how requires methods that handle those discrete interactions, yet few exist. Recently, it was discovered that even random networks exhibit dynamics that partitions the phase space into numerous attractor basins. Here we utilize this phenomenon to develop theory for the geometry of phase space partitioning in spiking neural circuits. We find basin boundaries structuring the phase space are pre-images of spike-time collision events. Formulating a statistical theory of spike-time collision events, we derive expressions for the rate of divergence of neighboring basins and for their size distribution. This theory reveals that the typical basin diameter grows with inhibitory coupling strength and shrinks with the rate of spike events. Our study provides an analytical and generalizable approach for dissecting how connectivity, coupling strength, single neuron dynamics and population activity shape the phase space geometry of spiking circuits.

---

Computing devices, whether natural or artificial, perform their function by finely orchestrated state changes of internal dynamical variables. In nervous systems these dynamical variables are physico-chemical states of nerve cells and synapses that connect them into complex networks called neural circuits. The causal dependencies arising from the synaptic interactions between cells greatly extend the space of functions computable by the circuit, beyond that of single neurons.

Mathematical models of neural circuits have been formulated in two fundamentally distinct ways ^1^. Most synaptic interactions in the brain are driven by sparsely-fired nerve impulses, called spikes, each lasting only a millisecond. In spiking neural network models this fundamental granularity of neuronal interactions is explicitly represented: all interactions depend on a discrete set of spike event times. Alternatively, continuous variable models for neural circuit dynamics are formulated by assuming that a frequency of nerve impulse generation, the firing rate, represents the information-encoding variable causally relevant for neural circuit computation. Firing rate models have been commonly used to model neural circuits ^2^, theoretically study their dynamics ^3,4^ and learning ^5–8^, and are the basis of spectacular advances in artificial computing systems ^9^. Statistical physics has played a role in this development, e.g. in clarifying the disordered phase space organization ^10^.

From a dynamical systems perspective, attractor states and their basins of attraction play a fundamental role in theories of neural computation. While analogous in some cases ^11,12^, however, rate models are not equivalent to temporally coarse-grained versions of spiking neural networks, even if they are closely matched in structure ^13^. Moreover, low firing rates (not much more than 1 Hz) in the cerebral cortex ^14^ make it hard to imagine how continuous rate variables associated to single neurons could provide a causally accurate description on behavioral time scales (hundreds of milliseconds). Developing theory for spiking networks may well require a dedicated approach. The absence of relevant averages and even a tractable ensemble of spiking trajectories, however, has thus far limited statistical approaches. Methods to design them ^15^ or to theoretically dissect the associated phase space organization are only starting to emerge.

Recently it has been discovered that, with dominant inhibition, even randomly wired networks partition their phase space into a complex set of basins of attraction, termed *flux tubes* ^16,17^. Here we utilize this setting to develop a statistical theory of phase space partitioning in spiking neural circuits. We first present a simulation study of flux tubes, uncovering their shape and revealing it is structured by a spike time collision event. Formulating these events, we then derive the conditions for and rate of the mutual divergence of neighboring tubes. Our main calculation is the derivation of the distribution of flux tube sizes, which we obtain from statistics of these events by leveraging the random connectivity to average over the disorder. Our analytical approach provides a transparent method to determine how coupling strength, connectivity, single neuron dynamics and population activity combine to shape the phase space geometry of spiking neural circuits.

## Methods

We study a tractable instance of the inhibition-dominated regime of neural circuits. *N* neurons are connected by an Erdős-Rényi graph with adjacency matrix *A* = (*A*_*mn*_). *A*_*mn*_ = 1 denotes a connection from neuron *n* to *m*, realized with probability, *p* = *K/N*. The neurons’ membrane potentials, *V*_*n*_ ∈(−*∞*, *V*_*T*_], are governed by Leaky Integrate-and-Fire (LIF) dynamics,

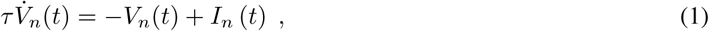

for *n* ∈ {1, …, *N*}. Here, *τ* is the membrane time constant and *I*_*n*_ (*t*) the synaptic current received by neuron *n*; when *V*_*n*_ reaches a threshold, *V*_*T*_ = 0, neuron *n* ‘spikes’, and *V*_*n*_ is reset to *V*_*R*_ = −1. At the spike time, *t*_*s*_, the spiking neuron, *n*_*s*_, delivers a current pulse of strength *J* to its *𝒪* (*K*) postsynaptic neurons, {*m |A*_*mns*_ = 1}, (*s* indexes the spikes in the observation window). The total synaptic current is

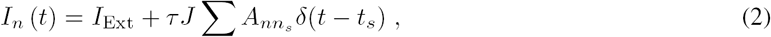

where *I*_Ext_ *>* 0 is a constant external current and *J <* 0 is the recurrent coupling strength. An 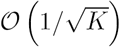-scaling of *J* is chosen to maintain finite current fluctuations at large *K* and implies that the external drive is balanced by the recurrent input. As a consequence, firing in this network is robustly asynchronous and irregular ^18–21^. Setting 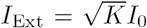, with *I*_0_ *>* 0, and 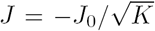 with *J*_0_ *>* 0, the corresponding stationary mean-field equation for the population-averaged firing rate, 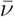, is ^17^

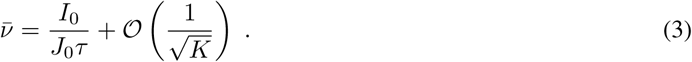

It is convenient to map the voltage dynamics to a pseudophase representation ^17,22^, 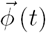, with

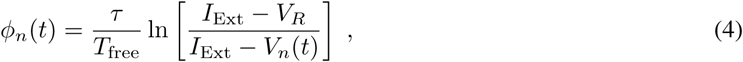

where *T*_*free*_ is the oscillation period of a neuron driven only by *I*_Ext_. *ϕ*_*n*_ (*t*) evolves linearly in time,

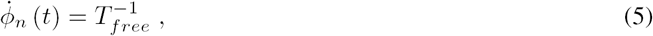

between spike events, *i.e. t* ∉ {*t*_*s*_}, and undergoes shifts given by the phase response curve, *Z*(*ϕ*), across input spike times where *ϕ* is the state at spike reception. In the large-*K* limit, *T*_free_ and *Z* (*ϕ*) simplify to

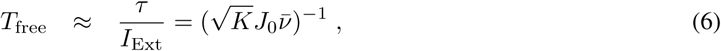

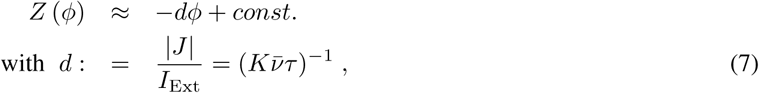

respectively (see Supplemental Methods for details). The differential phase response,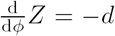, is essential for the strongly dissipative nature of the collective dynamics. For *J* = 0, the dynamics (equation (5)) would preserve phase space volume. This volume, however, is strongly contracted by spikes received in the post-synaptic neurons. Consider trajectories from a small ball of initial conditions as they emit the same future spike. The ball of phases at this spike contracts by a factor 1 − *d* along each of the *K* dimensions of the subspace spanned by the post-synaptic neurons. The volume thus contracts by (1 − *d*)^*K*^ → *e*^*λ*_inh_^ per spike, for *K* ≫ 1, with exponential rate,

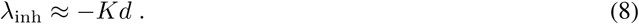

*λ*_inh_ *<* 0 is responsible for the linear stability of the dynamics given by equations (1) and (2), first shown in Refs. ^16,22^.

## Phase-space partitioning

The phase space volume taken up by an ensemble of nearby trajectories at a given spike is contracted at the spike’s reception. Larger phase space volumes, however, are not uniformly contracted but torn apart, with the pieces individually contracted and overall dispersed across the entire traversed phase space volume. The elementary phenomenon is illustrated in Fig. 1.

**Figure 1.**
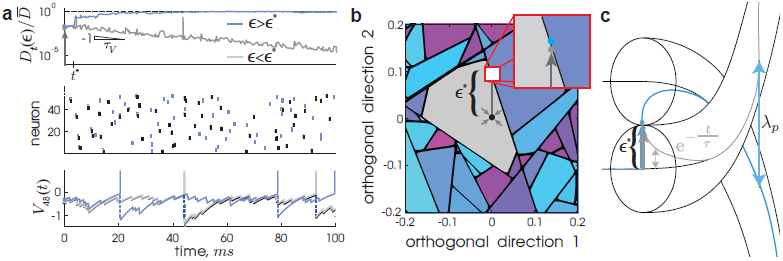
Finite-size perturbation instability and phase space partitioning in spiking networks. The three panels display the same slightly subcritical and supercritical perturbation of strength *∊*^*\**^*± δ*, *δ ≳* 0, respectively, applied once at *t* = 0 and in a random direction away from an equilibriated trajectory. **(a)** Temporal responses of the system. *Top*: The corresponding distance time series, *Dt*(*∊*), between the perturbed and unperturbed trajectories (gray: sub-critical, blue: super-critical; arrows in all three panels indicate the respective the perturbation). The divergence of *Dt*(*∊*^*\**^ + *δ*) begins at *t*^*\**^ ≈ 3 *ms*, and saturates at the average distance between randomly chosen trajectories, 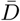 (dashed line) ^17^, while *Dt*(*∊*^*\**^− *δ*) only decays exponentially. *Middle*: The spike times as vertical ticks of the first 50 randomly labeled neurons from the network. The unperturbed sequence is shown in black. *Bottom*: The subthreshold voltage time course of an example neuron. The spike sequence and membrane potentials of the sub and supercritical trajectories decorrelate after *t*^*\**^. **(b)** A 2D cross-section (*δϕ*_1_, *δϕ*_2_) of the pseudophase representation of the phase space, orthogonal to and centered on the unperturbed trajectory from (a) at *t* = 0 (see also Ref. 15). The black dot at the origin indicates the latter, whose attractor basin is colored gray. The other colors distinguish basins in the local neighborhood. The two perturbed trajectories from (a) were initiated from (*δϕ*_1_, *δϕ*_2_) = (0, *∊*^*\**^*± δ*), respectively (shown as gray and blue dots, respectively, in the inset, in (a,Top and Bottom), and in (c)). **(c)** Schematic phase space caricature of two neighboring flux tubes with subcritical perturbations decaying on the order of the membrane time constant, *τ*, and typical basin diameter, *∊*^*\**^. The pseudoLyapunov exponent, *λ*_*p*_, is the rate at which neighboring tubes separate from each other (parameters: *N* = 200, *K* = 50, 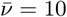, *τ* = 10 ms, *J*_0_ = 1).

We define the *critical perturbation strength*, *∊*^*\**^, as the flux tube’s extent out from a given state 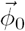 on the equilibriated trajectory, 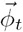, and in a given orthogonal perturbation direction, 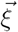,

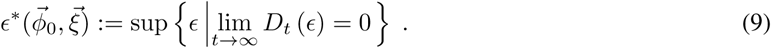

Here, *D*_*t*_ (*∊*) is the 1-norm distance,

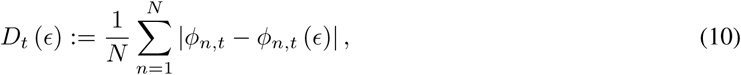

between 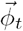 and the perturbed trajectory, 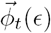 evolving freely from the perturbed state, 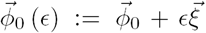 (reference time *t* = 0 and 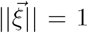; Supplemental Methods for details). *∊*^*\**^ is the largest value below which *D*_*t*_ (*∊*) vanishes in time. *D*_*t*_ initially decays exponentially, but for a supercritical perturbation, *∊* > *∊*^*\**^, there exists a *divergence event time, t*^*\**^ *>* 0, defined and obtained as the time at which a sustained divergence in *D*_*t*_ begins (see Fig. 1a).

A 2D cross-section of the phase space around 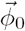 (Fig. 1b) reveals that the locations of these critical perturbations form lines which intersect to form polygon-shaped basin boundaries. Before developing a theory for this phase space organization (caricatured in Fig. 1c), we first analyze two main features of the geometry of a flux tube: the punctuated exponential decay of its cross-sectional volume and the exponential separation of neighboring tubes.

## Punctuated geometry of flux tubes

As expected from the typical phase space volume contraction (see equation (8)), we find along a simulated trajectory that the orthogonal phase space volume enclosed by the local flux tube exhibits exponential decay. This decay, however, is punctuated by blowup events. Figure 2a displays the spiking activity produced by the typical trajectory,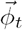 The neighborhood around 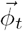 a time window is visualized in a *folded representation* using a fixed, 2D projection of the phase space (Fig. 2b and Supplementary Video; see Supplemental Information for construction details). The basin of attraction surrounding 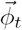 (Fig. 2b) consists of lines which remain fixed between spike times. Across spike times, new lines appear and existing lines disappear. At irregular intervals breaking up time windows of exponential contraction, large abrupt blowup events take the boundary away from the center trajectory (Fig. 2b, c), producing jumps in the area enclosed by the boundary. It is important to note that these events do not mean that the evolving phase space volume from an ensemble of states contained in the tube would expand. Such volumes only contract and converge to the same asymptotic trajectory. The basin of attraction itself, however, does not exclusively contract with time. In fact, it should on average maintain a typical size.

**Figure 2.**
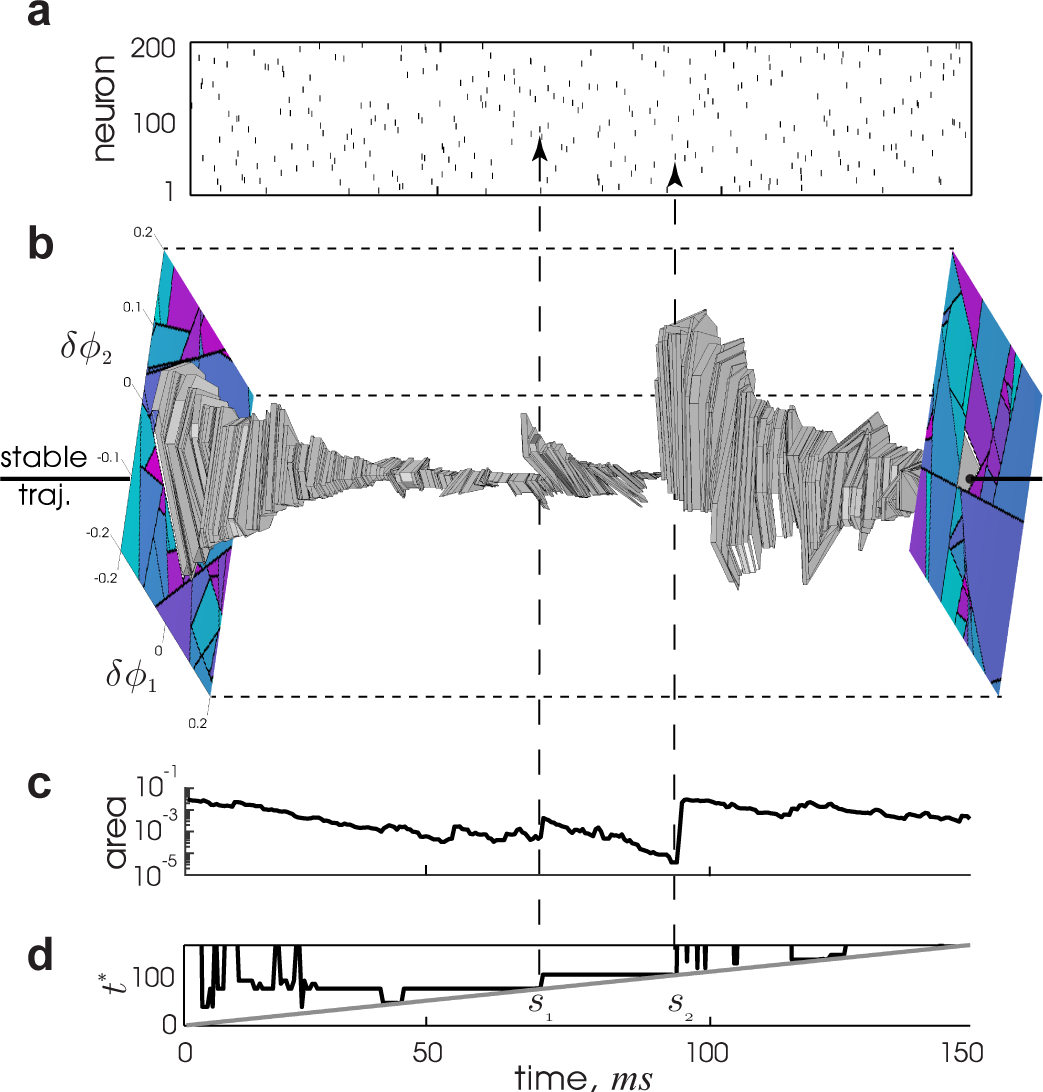
The basin boundary contracts towards and can blowup away from the stable trajectory within it. **(a)** Spike times from all neurons of the simulated trajectory,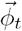, in a 150 ms window. **(b)** 2+1D folded phase space volume, (*δϕ*_1_, *δϕ*_2_, *t*), centered around 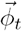 located at (0, 0, *t*) (black line) and extended in two fixed, random directions, 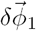 and 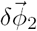. The center tube is filled gray in this volume, and the two cross-sections, (*δϕ*_1_, *δϕ*_2_, 0) and (*δϕ*_1_, *δϕ*_2_, 150), are shown. **(c)** Cross-sectional area of the center tube from (b) versus time. The area decays exponentially but can undergo abrupt expansions at blow-up times, e.g. at spikes *s*_1_ and *s*_2_ (note the logarithmic scale on the ordinate). **(d)** The absolute time of the next divergence event, *t*^*\**^ (see Fig. 1a, top), versus time, for perturbations along 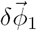. Note the step increase coincident with the blowup events seen in (b,c) (vertical, dashed lines). (Same parameters as Fig. 1.)

The blowup events typically coincide with a divergence event time, *t*^*\**^ (Fig. 1a), in some perturbation direction. Two such coincidences are visible in Fig. 2c,d. We conclude that the local basin at any time extends out in phase space until the perturbed trajectory approaches the pre-image of a divergence event occurring at a future time. Flux tube shape is then determined by the statistics of such events.

## Tube boundary and divergence

We analyzed a set of divergence events from simulations. We find that a collision of a pair of spikes constitutes the elementary event triggering the divergence of the perturbed trajectory. These pairs, hereon called *susceptible* spike pairs, were generated by connected pairs of neurons. Moreover, a perturbation-induced collision of a susceptible spike pair generated an abrupt spike time shift in one or both of the connected neurons’ spike times. We found that the nature of the spike time shift depends on the motif by which the two neurons connect. We denote the *backward-connected* pair motif *n*_*s*\*_ *← n*_*s*′_, where *s*^*\**^, the *decorrelation event index*, is the spike index of the earlier of the pair (note that *t*^*\**^ *≡ t*_*s*\*_), and *s*^′^ > *s*\* labels the later spike in the pair.

For *∊* ≲ *∊*\*, the presynaptic spike time, *t*_*s*′=*s*\*+1_, is advanced with increasing *∊* relative to the postsynaptic spike time, *t*_*s*\*_, until the two spikes collide (see Fig. 3). At collision (*∊* = *∊*\*), the pulsed inhibition and the rate of approach to voltage threshold cause an abrupt delay of *t*_*s*\*_ by Δ*t*_jump_. Using equation (3), we obtain

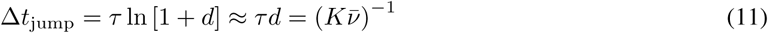

for *d* ≪ 1. Further details and the other two motifs (forward-connected and symmetrical) are discussed in the Supplementary Notes.

**Figure 3.**
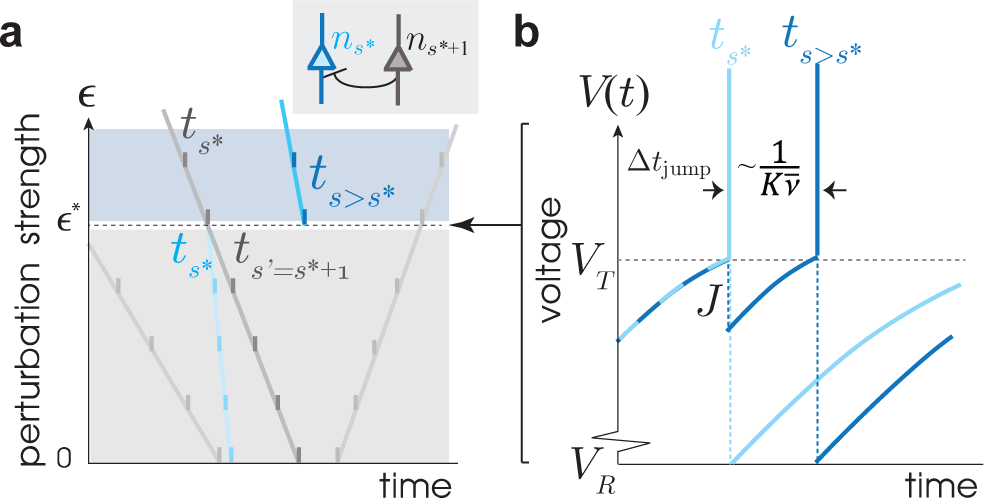
The collision of a susceptible spike pair causes an abrupt change in spike time. **(a)** A schematic illustration of the collision event for the backward-connected pair motif (shown in inset). For this motif, the interval vanishes as *∊* → *∊*^*\**^ from below. Perturbation strength, *∊*, is plotted versus time, where the timings of spikes for every perturbation strength are indicated as ticks on lines. The spike times shift continuously for *∊* < *∊*^*\**^. As the next input spike time, *t*_*s*\*^+1^_ (*∊*^*\**^ − *δ*), is advanced over *t*_*s*_*\** (*∊*^*\**^ + *δ*)A discontinuous jump of size Δ*t*_jump_occurs in the spike time of the post-synaptic neuron, *n*_*s*_*\** (light to dark blue) from *t*_*s*_*\** (*∊*^*\**^ − *δ*) to *t*_*s*′>*s*\*_ (*∊*^*\**^ + *δ*),*δ* ≿ 0. **(b)** Schematic illustration of the voltage of the *n*_*s*_*\** neuron versus time for *∊*^*\**^ *± δ*. The inhibitory kick of size 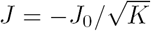 (not shown to scale) delays the spike time by an amount 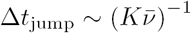.

For each spike in the network sequence, the rate of its susceptible spike partners is

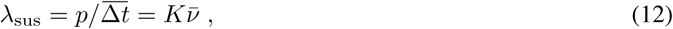

where 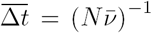 is the average distance between successive spikes. Since 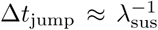, the spike time of neuron *n*_*s*\*_ is shifted forward typically as far as its next nearest susceptible partner spike. Thus, one collision event will typically induce another in at least one of the 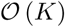 neurons to which the involved pair of neurons are presynaptic. A cascade of collision events then follows with near certainty (see Supplemental Notes for details).

The shift in *t*_*s*\*_ by Δ*t*_jump_ is carried forward to all future spike times of *n*_*s*\*_, so that *n*_*s*\*_ becomes a source of collision events. The total collision rate is then *λ*_sus_ multiplied by the number of source neurons, which approximately increments with each collision in the cascade. Averaging over realizations of the cascade (reference time *t*^*\**^ = 0), the average number of collisions, 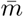, grows as 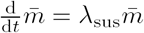. Finally, since each collision produces a jump in distance of equal size, we obtain the pseudoLyapunov exponent, *λ*_*p*_ = *λ*_sus_ from its implicit definition, 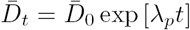 (see Supplemental Notes), as the exponential rate at which flux tubes diverge.

## Statistical theory of flux tube diameter

The geometry of a flux tube is captured by the *flux tube indicator function*, 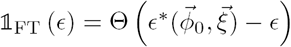, evaluated across network states,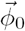, of its contained attracting trajectory and perturbation directions,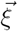. Using the Heaviside function, Θ(*x*), 𝕝_FT_ (*∊*) = 1 for perturbations remaining in the tube (*∊* < *∊*^*\**^), and 0 otherwise. The average of 𝕝^FT^ (*∊*) over 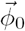 and 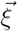,

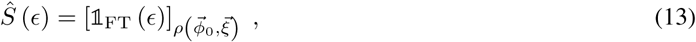

is the *survival function*: the probability that an *∊*-sized perturbation does not lead to a divergence event later in the perturbed trajectory, *i.e. ∊* < *∊*\*, and is formally defined as 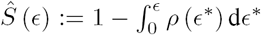, with *ρ* (*∊*\*) the transformed density over *∊*\*. 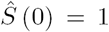 and decays to 0 as *∊ →∞*. The scale of this decay defines the typical flux tube size. Calculating 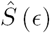 requires two steps: firstly, establishing a tractable representation of 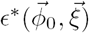 and secondly, performing the average in equation (13). Both of these in general pose intricate problems. However, as we will see next, both substantially simplify when generic properties of the asynchronous, irregular state are taken into account.

Perturbed spike intervals are obtained using the spike time deviations, *δt*_*s*_ (*∊*) := *t*_*s*_ (*∊*) − *t*_*s*_ (0), *s* = 1, 2, *…*,

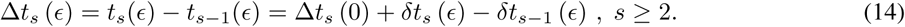

In a linear approximation we find,

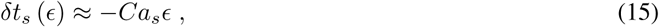

where 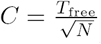 converts network phase deviation to spike time deviation and *a*_*s*_ is a dimensionless susceptibility that depends on the adjacency matrix, *A* = (*A*_*mn*_), derivatives of the phase response curve evaluated at the network states at past spike times, 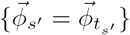 for *s*′ < *s*, and the perturbation direction 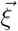 (see Supplemental Notes for its derivation). Substituting equation (15) into equation (14) gives

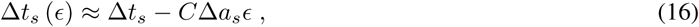

with Δ*t*_*s*_ = Δ*t*_*s*_ (0). Note that Δ*t*_*s*_ (*∊*) can have a zero, *i.e.* a spike time collision only when Δ*a*_*s*_ = *a*_*s*_ −*a*_*s*−1_ *>* 0.

To obtain the scaling behavior of the flux tube geometry it is sufficient to examine the statistics of flux tube borders using the corresponding divergence events generated by collisions of backward-connected susceptible spike pairs in the perturbed trajectory (Fig. 3). In these cases, the perturbation strength *∊* → *∊*\* as the network spike interval 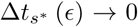 for 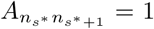. In fact, the latter condition serves in these cases as an implicit definition of *∊*\* and *s*\*.

According to equation (16), 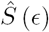 in principle depends on the adjacency matrix, *A* = (*A*_*mn*_), of the network realization. Removing this dependence by averaging over the ensemble of graphs, *P*_*A*_ ((*A*_*mn*_)), simplifies the calculation of the survival function,

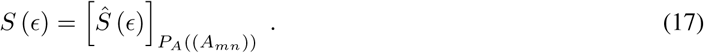

Evaluating the right-hand side of equation (17) using the perturbed spike intervals, linearized in *∊*, requires knowledge of the joint probability density of all variables present in equation (16),

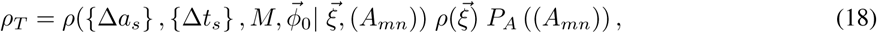

where we have chosen the perturbation direction, 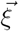, to be statistically independent of the state,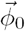, being perturbed at *t* = 0. Here, the unperturbed spike pattern is represented by two random variables: *M*, the number of spikes in the time interval [0, *T*] after the perturbation, and {Δ*t*_*s*_}, the set of all *M* − 1 inter-spike intervals in this window. It is well understood that in the large-system limit in a sparse graph, 1 ≪ *K* ≪ *N*, the currents driving individual neurons in the network converge to independent, stationary Gaussian random functions ^23^. For low average firing rates, this implies that the pattern of network spikes (*M*, {Δ*t*_*s*_}) resembles a Poisson process ^24^. Furthermore, the susceptibility becomes state-independent in this limit. Neglecting the weak dependence between the distribution of network spike patterns and *A* = (*A*_*mn*_), the full density, *ρT* (equation (18)), approximately factorizes,

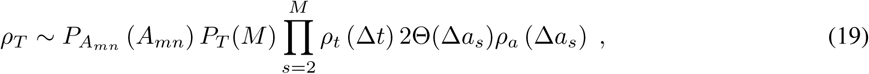

with distribution of a single adjacency matrix element, *P*_*A*_*mn*__ (*A*_*mn*_ = 1) = *p*, *P*_*A*_*mn*__ (*A*_*mn*_ = 0) = 1 − *p*, count distribution of spikes in the observation window, *P*_*T*_ (*M*), and distribution of single inter-spike interval *ρ*_*t*_ (Δ*t*). The latter is exponential with rate 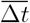. With these approximations (see Supplementary Notes for details), all dependencies on the distribution of perturbation direction are mediated by the susceptibilities, {Δ*a*_*s*_}. For any isotropic 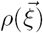 having finite-variance, *ρ*_*a*_ (Δ*a*_*s*_) has zero mean and standard deviation proportional to 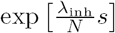, with the average contraction rate per neuron, 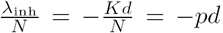, due to the inhibition. The factor 2Θ(Δ*a*_*s*_) places support only positive values of Δ*a*_*s*_ as required.

As *ρ*_*T*_ factorizes, so does *S* (*∊*),

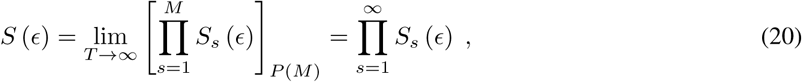

where *S*_*s*_ (*∊*) is the probability that a perturbation of strength *∊* does not lead to a collision event involving the *s*^th^ spike. With the above simplifications,

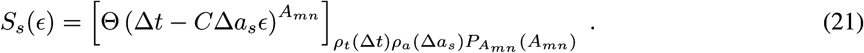

Evaluating equation (21) (see Supplementary Notes for the derivation), we find

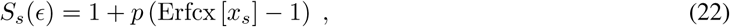

where 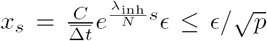, and Erfcx 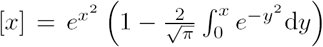 is the scaled complementary error function. Erfcx [*x*_*s*_] − 1 ≈ −*x*_*s*_ for 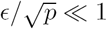, so that finally

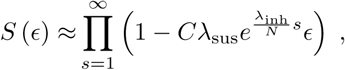

where we have identified 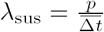. Employing the logarithm and 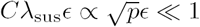,

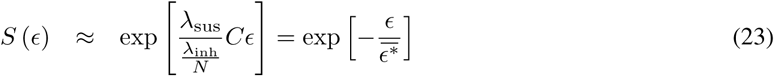

with

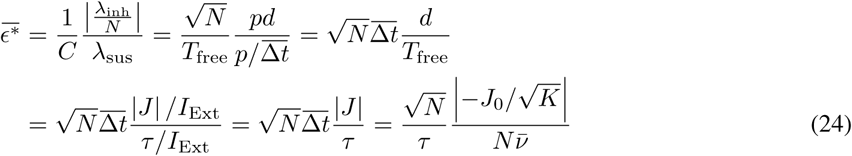

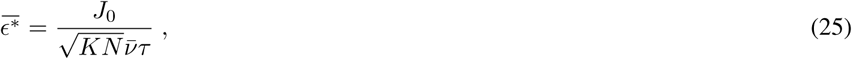

where we have used equations (6) and (7) in the second line and note the cancellation of *p* and *I*_Ext_. Equation (23) shows for 1 ≪ *K* ≪ *N* that the basin diameter, *∊*\*, is exponentially distributed and so completely determined by its characteristic scale, 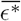 (equation (25)), that is smaller for larger network size, higher average in-degree, higher population activity, and larger membrane time constant, *τ*. 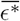 grows, however, with the synaptic coupling strength, *J*_0_. In Fig. 4b, we see quantitative agreement in simulations between the definition of 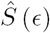 (equation (13) using the definition of *∊*\*, equation (9)) and its approximate microstate parametrization (equations (20), (21)). These also confirm the exponential form of our reduced expression (equations (23), (25)) and a scaling dependence on *J*_0_ (Fig. 4c). The latter holds until *J* is no longer of size 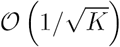. The other scalings were reported in Ref. ^17^. A derivation of only the characteristic scaling of 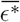, but not depending on the Poisson spiking assumption, is given in the Supplemental Notes.

**Figure 4.**
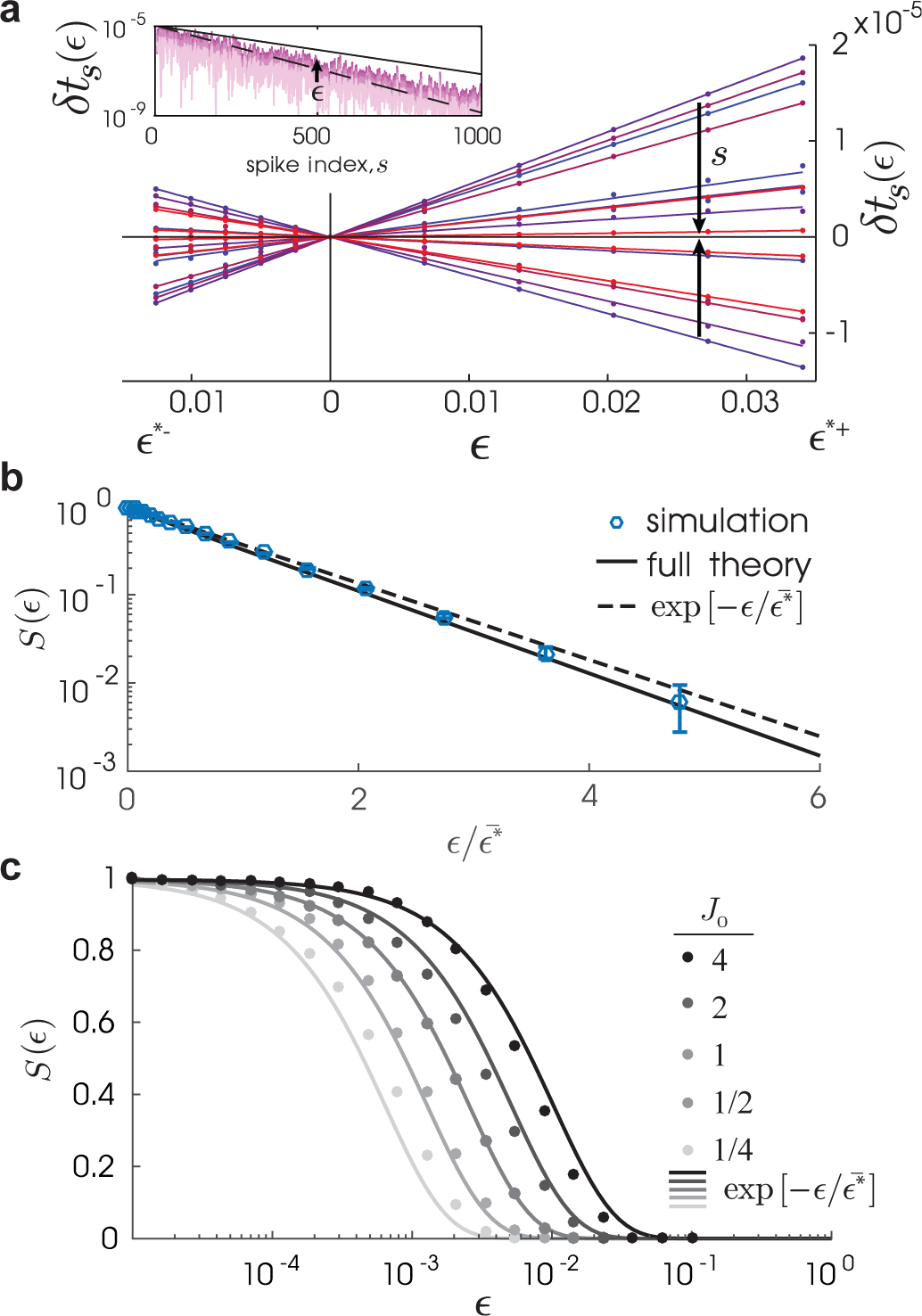
The flux tube indicator function, once expressed with microstate variables and averaged, gives the survival probability to remain in the containing flux tube. **(a)** Spike-time deviations, *δt*_*s*_ (*∊*) (dots), as a function of perturbation strength up to the positive and negative critical strength, *∊*^*\*−*^ and *∊*^*+^, respectively, for *s* = 1, *…,* 15 (colors) with their linear approximation (lines) given by equation (15). Inset: *δt*_*s*_ (*∊*) as a function of *s* (shown for *∊* = 0.2*∊*^*\*±*^, 0.4*∊*^*\*±*^, 0.6*∊*^*\*±*^, 0.8*∊*^*\*±*^) decays exponentially at a rate near the maximum and mean Lyapunov exponent, *λ*_max_ (black line) and *λ*_mean_ (black-dashed line) respectively ^17^. **(b)** The survival probability function *S* (*∊*) from simulations (dots, equation (13); bars are standard error), theory (line, equations (20),(21)), and the simplified theory at large *K*, 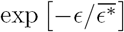 (dotted line, equation (23)), where 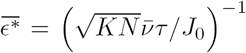. **(c)** *S* (*∊*) from simulations (dots) and 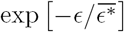 (lines) for *J*_0_ = 2^*n*^, *n* = −2, −1, 0, 1, 2. (Same parameters as Fig. 1 except *N* = 10^4^, *K* = 10^3^.)

## The geometry of phase space partitioning

Figure 5 presents the phase space organization of these spiking circuits as we have revealed it, replacing the caricature of Fig. 1c. For a perturbation made to a stable trajectory, the geometry of the determining collision event is shown in Figure 5a, in a folded representation. The pre-images of this event determine the flux tube boundary back to the perturbation. Our results also provide a global, i.e. non-folded geometry of the partitioning (Fig. 5b(left)). Susceptible spike collisions are edges of the *N*-dimensional unit hypercube of phases where the corresponding voltages of two connected neurons both approach threshold. The Poincare section obtained by projecting the dynamics orthogonal to the trajectory (since no motion exists orthogonal to this subspace) then reveals the intrinsic partition. Here, the polygon basin boundaries arise as the pre-images of the projections of susceptible edges lying nearby the trajectory at future spike times (Fig. 5b(right)).

**Figure 5.**
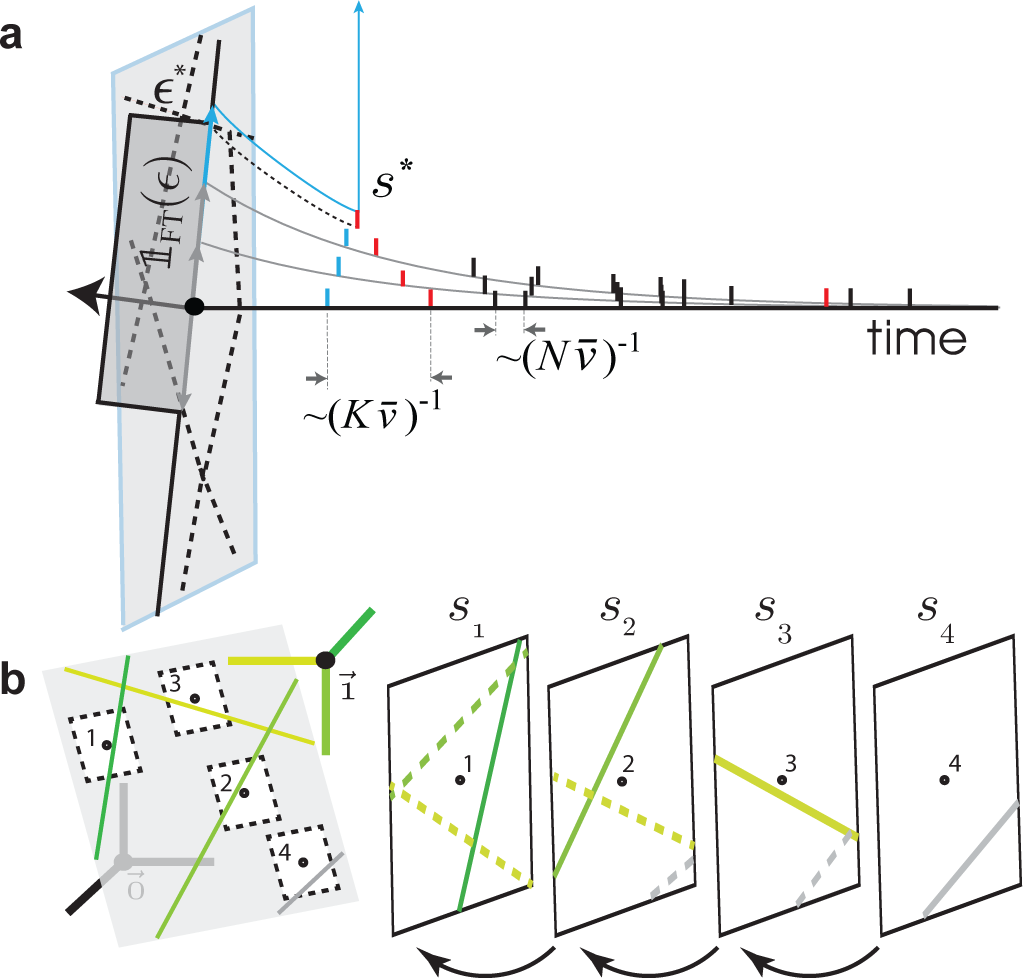
Flux tube boundaries are the pre-images of future susceptible spike collisions. **(a)** A folded phase space representation of a susceptible spike collision. Spikes (ticks) occur at a rate 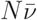 in the unperturbed trajectory (black line). For an example spike (blue tick), its susceptible spike partners (red ticks) occur at lower rate 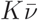. Small perturbations (gray arrows) lead to trajectories (gray) exhibiting spike time deviations that decay over time (tick alignment). A larger perturbation (example just beyond the critical perturbation strength: blue arrow) can push a spike and one of its susceptible partner spikes in the subsequent trajectory (blue and red, respectively) to collide, generating a divergence event at spike *s*\*. The indicator function, 𝕝_FT_ (*∊*), has support (dark gray) only over the local tube. **(b)** Constructing the local flux-tube partition in the non-folded phase space. *Left*: Susceptible spikes are represented by susceptible edges (thick green lines) of the unit hypercube having 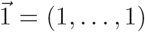 (black dot) as an endpoint. An intrinsic random partition (thin green lines) is generated by projecting these edges onto the hyper-plane orthogonal to 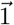 (light gray). A given trajectory (labeled sequence of small dots) and its local neighborhood (within black dashed lines) is shown. *Right*: The flux tube partition for this trajectory at a given spike (here *s*_1_) is obtained from back-iterating the intrinsic partition from all future spikes (here only partitions from *s*_2_, *s*_3_, and *s*_4_ are back-iterated; dashed lines). The partition at sufficiently distant future spikes (here the gray edge at *s*_4_) will no longer refine the partition in the local neighborhood at *s*_1_, since the expansive backwards dynamics maps the projected edges outside the neighborhood. A concrete example obtained from simulations is presented in the Supplemental Notes.

## Discussion

We have developed a theory of phase space partitioning in spiking neural circuits, exemplified using the phenomenon of flux tubes. Importantly, the approach yields the dependence on various control parameters. We find the flux tube diameter contracts with the rate of volume contraction per neuron, 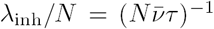, due to the inhibition received across the post-synaptic subspace of each spike. This contraction is punctuated, however, by collision events between susceptible spikes, *i.e.* those from pairs of connected neurons, occurring at rate 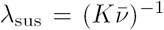 and across which the basin volume expands out to a pre-image of the next collision event. For some neighboring tube, this collision event sets off a cascade of such events with exponential rate, *λ*_sus_ that is responsible for their mutual divergence. Using these collision events to identify the spiking trajectories lying on flux tube boundaries, we were able calculate the size distribution of these basins. The average size is controlled by the ratio of these two exponential rates. Leaving out a factor converting shifts in spike time to shifts in state,

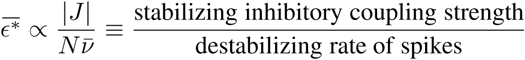

The final scaling,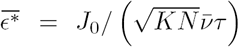, thus combines the contraction from the single neuron dynamics responsible for the dissipative dynamics, with the overall rate of spikes, which appears since each spike can be involved in a destabilizing collision event. Both contracting and expanding rates scale with the probability of connection, *p*, so we intuitively expect *p* to appear in 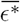 only implicitly through *J* and, reassuringly, *p* indeed cancels out.

Our framework motivates a variety of extensions. Our calculations can be performed for different disordered connectivity ensembles (e.g. correlated entries from annealed dilution processes ^25^ and structured second-order statistics ^26^), different activity regimes (e.g. non-Markovian spike interval processes ^27^), and different single neuron models (e.g. any threshold neuron with known phase response curve). We have applied the theory to an instability caused by abrupt changes in spike time due to an inhibitory input near voltage threshold, a scenario that can also be analyzed in neuron models with smooth thresholds (e.g. the rapid theta-neuron ^28^ that has the LIF neuron as a limit). The theory may also apply to other, as yet unknown instabilities involving spike collision events. Finally, while the linear stability of the dynamics precludes finite, asymptotic (Kolmogorov-Sinai) entropy production, the partition refinement picture we provide in Fig. 5b suggests a transient production of information about the perturbation on timescales of the order of the divergence event time, *t*\*. Making this connection to ergodic theory more precise is an interesting direction for future research.

Applying our approach in a relatively idealized context allowed for a tractable assessment of phase space organization. Despite its simplicity, however, the LIF neuron accurately captures many properties of cortical neurons, such as their dynamic response ^29^. We have also neglected heterogeneity in many properties. For instance, in contrast to the locally stable regime studied here, mixed networks of excitatory and inhibitory neurons can instead be conventionally chaotic ^30^. This chaos can nevertheless be suppressed in the ubiquitous presence of fluctuating external drive ^31,32^ or with spatially-structured connectivity ^33^, suggesting a generality to locally stable dynamics and phase space partitioning in neural computation. Our approach, in particular the way we have quantified the ensemble of perturbed spiking trajectories, can inform formulations of local stability in these more elaborate contexts. Of particular interest are extensions where a macroscopic fraction of tubes remain large enough to realize encoding schemes tolerant of intrinsic and stimulus noise. For example, extensions to random dynamical systems ^34,35^ could provide theoretical control over spiking dynamic variants of rate network-based learning schemes to generate stable, input-specific trajectories ^7^.

Recent advances in experimental neuroscience have allowed for probes of the finite-size stability properties of cortical circuit dynamics call for *in vivo*. For example, simultaneous intra- and extra-cellular recordings in the whisker motion-sensing system of the rat reveal that the addition of a single spike makes a measurable impact on the underlying spiking dynamics of the local cortical area ^36^. Indeed, rats can be trained to detect perturbations to single spikes emitted in this area ^37^. Representative toy theories, such as the one we provide, can guide this work by highlighting the features of spiking neural circuits that contribute to phase space partitioning. The combined effort promises to elucidate the dynamical substrate for neural computation at the level at which the neuronal interactions actually operate.

## Acknowledgements

M.P.T. would like to acknowledge discussions with Michael Monteforte, Sven Jahnke and Rainer Engelken. This work was supported by BMBF (01GQ07113, 01GQ0811, 01GQ0922, 01GQ1005B), GIF (906-17.1/2006), DFG (SFB 889), VW-Stiftung (ZN2632), and the Max Planck Society.

## Author Contributions

M.P.T. conceived the project, developed the concepts, and wrote the manuscript. F.W. supervised the project, discussed the results and edited the manuscript.

## Additional Information

The authors declare no competing financial interests.

